# Exploring soil microbes’ potential to assess Atlantic Forest restoration trajectories

**DOI:** 10.64898/2026.09.11.747570

**Authors:** Haroldo Borges Gomes, Pedro M. Pedro, Thomas Püttker, João Francisco Coelho, Laury Cullen, Marcelo Rodrigo Alves

## Abstract

Soil microbe communities are key indicators of ecosystem recovery, yet their integration into operational restoration monitoring remains limited. We evaluated the potential for benchmarking early Atlantic Forest restoration with reference site-based comparisons using bacteria and fungi. Samples were collected from three geographic clusters and spanned a pasture-restoration-forest land-use gradient. Using metabarcoding, we assessed whether soil microbe community diversity, structure, and functional profiles respond to restoration progress considering influences from spatial arrangement and/or successional stage (determined using NDVI). We found that OTU alpha diversity was not explained by either predictor, whereas the Shannon diversity of bacterial functions declined significantly with increasing NDVI. Bacterial and fungal OTU composition (Bray-Curtis) responded significantly to both NDVI and geographic cluster, as did microbe functional profiles, although only NDVI proved both marginally and conditionally significant. We also found that soil chemical parameters were primarily structured by geographic cluster but not by NDVI, suggesting that many abiotic edaphic conditions reflect soil history rather than current vegetation status. These findings suggest that both microbe taxonomic and functional metrics track revegetation progress and validate eDNA as a scalable, sensitive restoration monitoring framework. Importantly, our results also support the use of reference site-based monitoring for Atlantic Forest restoration.

## Introduction

Large-scale ecological restoration has emerged as an important strategy to address high levels of atmospheric carbon, biodiversity loss and degradation of ecosystem functions. In this context, ambitious restoration targets have been established for the Brazilian Atlantic Forest under national and international commitments (Crouzeilles et al., 2019). However, the success of these initiatives depends not only on the net area restored, but on the ecological directionality and pace of recovery. Also relevant is that sponsors of both small- and large-scale restoration initiatives now increasingly require reporting of ecological metrics.

To fulfill restoration’s ecological goals, the biotic indicators monitored in restored areas and their ecological functions must gradually approximate those of the original habitats. This has come to be known as restoration’s “ecological trajectory”. As such, current restoration assessments reframe outcomes not as a binary success/failure but as a goals-based framework toward a predefined reference state, typically a nearby remnant ecosystem. The Society for Ecological Restoration’s 5-Star Recovery System, for example, provides a tiered framework for evaluating the degree of recovery of a restored site relative to such a reference (Gann et al., 2019; Campbell et al., 2024).

Restoration monitoring has historically relied on aboveground metrics such as vegetation structure and animal assemblages. While valuable, these indicators offer limited insight into the belowground recovery processes that underpin long-term resilience and functioning of terrestrial ecosystems (Bardgett & Van Der Putten, 2014). Soil microbe communities inform a broad swath of this ecological dimension, since they govern important ecological functions including nutrient cycling, organic matter decomposition, plant-soil feedback, and pathogen regulation, among others (Delgado-Baquerizo et al., 2016). Consequently, soil microbes are powerful bioindicators of early recovery and appropriate metrics for adaptive management of restored areas (Harris, 2009).

There is growing recognition that next-generation monitoring frameworks that leverage soil environmental DNA (eDNA) can deliver sensitive, scalable, actionable and functionally meaningful assessments of ecological status (Banning et al., 2011; Liddicoat et al., 2022). Applying these frameworks to restoration trajectories necessarily requires understanding the impact of geographic distance to the reference site and how changes in soil properties (especially those independent of the restoration process) shape microbe communities. Ideally, i) microbial turnover during succession should not be strongly confounded by geographic distance between the restored and reference sites, particularly because reference sites are often distant to restored sites; and ii) microbial community turnover should reflect the restoration gradient, i.e. restored sites should host microbiomes intermediate between the original matrix (often pasture in the Atlantic Forest) and regional reference forests.

We sought to evaluate if eDNA provides a useful metric for the operational monitoring of restoration success in the Atlantic Forest. By benchmarking restored sites against native forest targets in the Pontal do Paranapanema region of São Paulo state, this study assesses the relative importance of geographic location, vegetation density (NDVI), and soil chemical properties for explaining the structuring of microbial OTU and functional communities across a land-cover gradient from pasture (the usual restoration baseline in the region), to restored sites to mature native forest reference sites.

## Methods and materials

### Study design and sample collection

We sampled soil from 12-24 June 2023 within three geographic clusters (subsequently *clusters I*, *II*, and *III*). Soil from *clusters I* and *II* represented three land cover types; each had samples categorized as pasture (n=3), restoration (>15 or >10 years; n=3) and forest (reference; n=3). *Cluster III* contained only soil from restoration (<3 years, n=3) and reference forest (n=3), as no viable pasture site was available here for comparison (Supplementary file *Sites.kml*).

Each sampling site was delineated by a one-hectare perimeter and, within this, 10 soil samples were randomly collected to a depth of 20-cm using a Dutch-type auger.

Because many of these samples were collected from triads in close spatial proximity (Supplementary file *Sites.kml*) and often shared land-use history, we considered samples from the same cluster and land use as spatially dependent and therefore used mean values of adjacent samples in all analyses to avoid pseudo-replication prior to downstream analyses. This yielded a total of eight samples for statistical analyses.

### NDVI calculation

Given the range of restoration ages represented at the restored sites (<3, >10 or >15 years), NDVI was used as a proxy for vegetation recovery, capturing differences in density and canopy development across pastures, restored sites, and forest areas. Mean monthly NDVI values were calculated for each sampling site for May, June and July 2023 using the COPERNICUS/S2_SR_HARMONIZED Sentinel-2 surface reflectance collection in the Google Earth Engine.

### Soil Chemical Analyses

For each sample, a portion of the soil collected was assessed for 41 chemical parameters by a commercial provider. These encompassed the common agricultural parameters listed in *SupplementalData.xlsx*, TAB “*ChemicalMetrics*”.

### Molecular Analysis

Total genomic DNA was extracted from 250-mg of soil according to manufacturer’s instructions using either of two kits: DNeasy PowerSoil (Qiagen) or E.Z.N.A. Soil DNA (Omega). Bacterial and fungal markers were used to generate sequence-based microbial profiles via Illumina MiSeq sequencing for 2x250-bp paired-ends reads. Bacteria PCRs targeted the 16S rRNA V3-V4 operon with primers 341F (5’-CCTAYGGGRBGCASCAG-3’) and 806R (5’-GGACTACNNGGGTATCTAAT-3’) (Takahashi et al., 2014). Fungal ITS1 PCRs used primers ITS5-1737F (5’-GGAAGTAAAAGTCGTAACAAGG-3’) and ITS2-2043R (5’-GCTGCGTTCTTCATCGATGC-3’) (White et al., 1990).

### Bioinformatics

Bioinformatic sequence processing used *MOTHUR* v.1.36.1 (Schloss et al., 2009) to filter Illumina sequences with a minimum average quality score of 25. We allowed for no nucleotide differences in the barcode region of the oligo and four differences in the priming region. Clustering of reads into OTUs was as in the USEARCH UPARSE pipeline (https://www.drive5.com/usearch/manual/uparse_pipeline.html). A read-clustering threshold of 3% was adopted to bin OTUs.

Taxonomic assignment of filtered bacterial and fungal OTUs was done with the DADA2 pipeline in R (R Core Team, 2023) using the *assignTaxonomy* function (Callahan et al., 2016) against the SILVA reference database for bacteria (v138.2; Quast et al., 2013) and the UNITE dataset release v.19.02.2025 (Abarenkov et al., 2024).

### Functional guilds

Bacterial functional guilds were predicted using PICRUSt2 v. 2.6.3 (Douglas et al., 2020) and encompassed pathway abundance predictions from enzyme commission (EC) annotations. Fungal functional guilds were assigned via the *fungaltraits* R package (Põlme et al., 2020) for genus-level assignments exclusively, as there were relatively few species-level OTUs. Assignments to compound functional guilds, that is, those genera assigned to more than one fungal guild, were removed from the analyses to not confound functional profiles.

### Statistical analyses

#### Biotic profiles (OTU- and function-based analyses)

Possible effects of *cluster* and *NDVI* on variation in Shannon Diversity (alpha diversity) from i) bacterial OTUs, ii) bacterial functional groups, iii) fungal OTUs and iv) fungal functional guilds were partitioned using the *varpart* function of the *vegan* r package. This assessed both the marginal and conditional contributions of geography (*cluster* identity) and vegetation density (*NDVI*). Significance was assessed by fitting the marginal, conditional (using *Condition*()), and full *NDVI*+*cluster* models with *rda*() on the Shannon response and testing each with *anova.cca*() with 999 permutations.

The variation in community composition among sampling sites (beta diversity) was visualized with NMDS based on Bray-Curtis dissimilarity in *vegan* with e*nvfit* used to test the fit of *NDVI* and *cluster* values onto the NMDS axes (Oksanen et al., 2020). PERMANOVA was used to test the significance of marginal and conditional contributions from *cluster* identity and *NDVI* to community composition.

Spearman correlation assessed the relationship between fungal and bacterial Shannon diversity (for both OTU and function). Finally, in order to assess whether the bacteria and fungi beta diversity showed correlated patterns across samples, we utilized Mantel and Partial Mantel tests of their Bray-Curtis distance matrices (for both OTU and functional distances).

#### Abiotic profiles (soil chemistry)

We undertook an ordination of sample chemical profiles with principal components analysis (PCA) using the R *prcomp* function with centered and scaled data on standardized soil variables (n=41).

As was done for Shannon diversity above, we assessed how soil chemistry varied in relation to geography *(cluster identity) and* vegetation density *(NDVI)*. (We did this separately from the analogous alpha diversity calculations above because the small sample sizes precluded analysis with more than two explanatory variables). Possible effects of the explanatory variables on soil chemistry variation (the full standardized 41-variable chemistry matrix) were partitioned using the *varpart*() function in *vegan*. Significance was assessed by fitting the marginal, conditional (using *Condition*()), and full *NDVI*+*cluster* models with *rda*() on the chemistry response and testing each with 999 permutations in *anova*.*cca*().

All statistical analyses employed the *vegan* package in R v.4.3.1 unless otherwise noted.

## Results

### Microbe sequence results

After sequence filtering, bacteria were represented by 9,356 and fungi by 7,339 unique OTUs (at 97% similarity). PICRUSt2 functional assignments of these OTUs were successful for all but one bacterial OTU and predicted 424 bacterial metabolic pathways. Fifteen percent (1,106) of fungal OTUs could be assigned to one of 16 unambiguous guilds, dominated by *plant pathogens*, *undefined saprotrophs*, *endophytes*, and *animal pathogens*. Microbial phylum and functional composition for all samples is shown in *SupplementalData.xlsx,* TAB “*taxonomic_composition”*.

### Microbial Shannon alpha diversity

Neither bacterial nor fungal Shannon OTU diversity was significantly explained by *NDVI*, *cluster* identity, or their combinations (Table 1). The variance in Shannon functional diversity of bacteria, however, was significantly negatively correlated to both marginal and conditional *NDVI*. Interestingly, fungal functional diversity varied significantly only among *clusters*, indicating a localized effect where *cluster I* samples had higher Shannon diversity values than the other two (Table 1; Figure 1).

**Table 1:** Variance partitioning (*varpart*) results of microbial OTU-based and functional-based Shannon diversity. The variation in diversity values was tested against *NDVI* and *cluster* predictors. Significant associations are in bold/underlined

| BACTERIA OTU<br>SHANNON DIVERSITY | Source | Adj. R<sup>2</sup> | F | p-value | FUNGAL OTU<br>SHANNON DIVERSITY | Source | Adj. R<sup>2</sup> | F | p-value |
| --- | --- | --- | --- | --- | --- | --- | --- | --- | --- |
|  | NDVI + cluster | 0.022 | 1.05 | 0.472 |  | NDVI + cluster | 0.470 | 3.07 | 0.175 |
|  | NDVI | -0.025 | 0.83 | 0.404 |  | NDVI | 0.106 | 1.83 | 0.215 |
|  | cluster | 0.058 | 1.22 | 0.378 |  | cluster | 0.230 | 2.04 | 0.236 |
|  | NDVI\| cluster | -0.036 | 0.82 | 0.412 |  | NDVI \| cluster | 0.240 | 3.26 | 0.190 |
|  | cluster \| NDVI | 0.048 | 1.15 | 0.397 |  | cluster \| NDVI | 0.364 | 3.06 | 0.157 |
|  | shared | 0.011 | — | — |  | shared | -0.134 | — | — |
|  | residuals | 0.978 | — | — |  | residuals | 0.530 | — | — |
| BACTERIA<br>FUNCTIONAL SHANNON<br>DIVERSITY | Source | Adj. R<sup>2</sup> | F | p-value | FUNGAL<br>FUNCTIONAL SHANNON<br>DIVERSITY | Source | Adj. R<sup>2</sup> | F | p-value |
|  | <u>NDVI + cluster</u> | <u>0.723</u> | <u>7.09</u> | <u>0.045</u> |  | NDVI + cluster | 0.517 | 3.50 | 0.124 |
|  | <u>NDVI</u> | <u>0.749</u> | <u>21.93</u> | <u>0.001</u> |  | NDVI | -0.166 | 0.002 | 0.969 |
|  | cluster | -0.326 | 0.14 | 0.891 |  | <u>cluster</u> | <u>0.613</u> | <u>6.544</u> | <u>0.049</u> |
|  | <u>NDVI \| cluster</u> | <u>1.049</u> | <u>19.93</u> | <u>0.008</u> |  | NDVI \| cluster | -0.096 | 0.009 | 0.942 |
|  | cluster \| NDVI | -0.027 | 0.71 | 0.530 |  | cluster \| NDVI | 0.684 | 5.249 | 0.076 |
|  | shared | -0.300 | — | — |  | shared | -0.071 | — | — |
|  | residuals | 0.277 | — | — |  | residuals | 0.483 | — | — |

**Figure 1:**
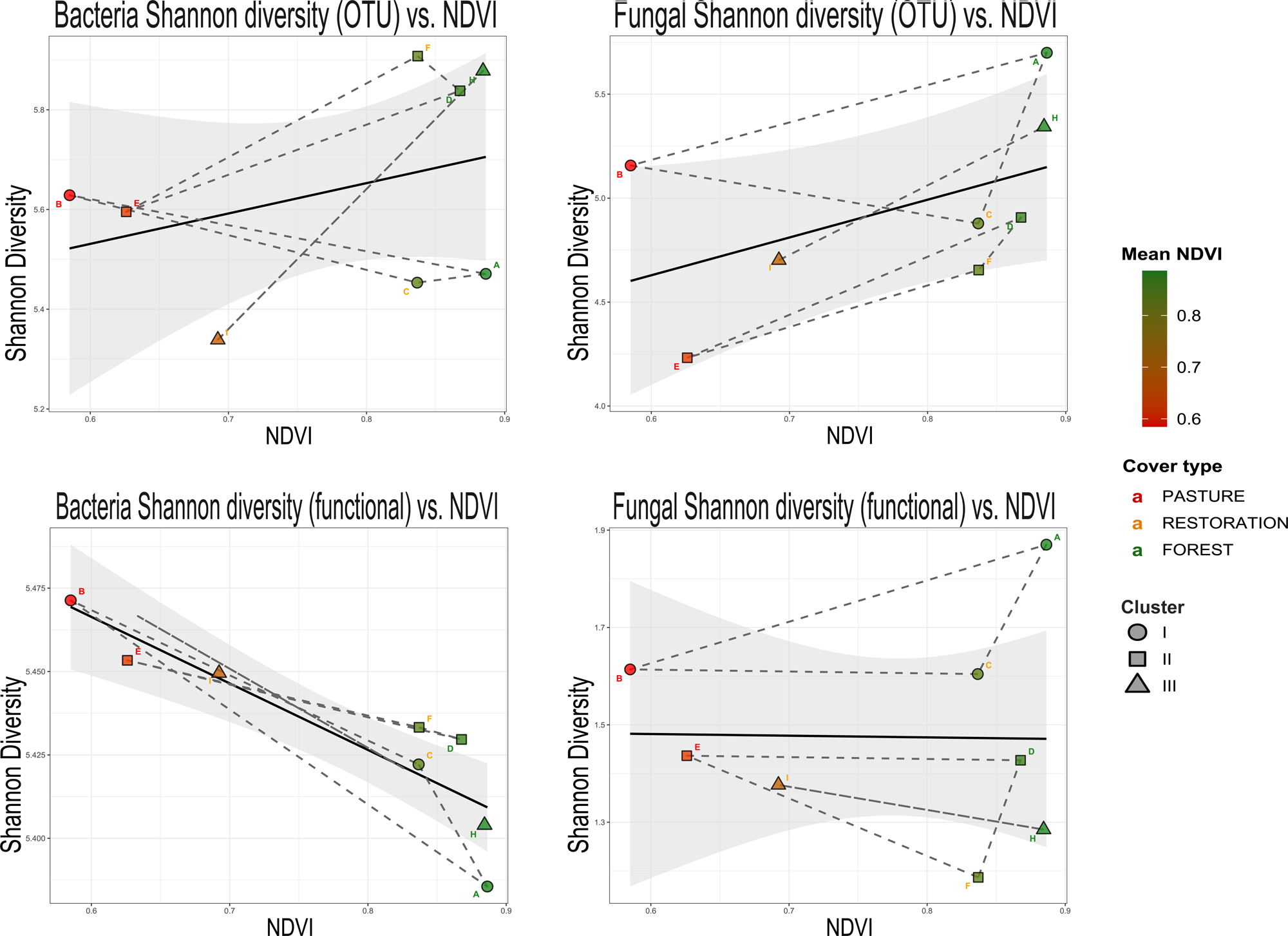
Shannon diversity of OTU (top-row) and functional (bottom row) values against *NDVI* for bacteria (left) and fungi (right). Dashed lines connect samples from the same geographic *cluster*.

### Microbe beta diversity

For both microbe communities, NMDS showed a clear segregation of samples based on *NDVI* and *cluster* values for both OTU or function datasets. However, the *envfit* estimate of ordination space was only significant for NDVI (Figure 2).

**Figure 2:**
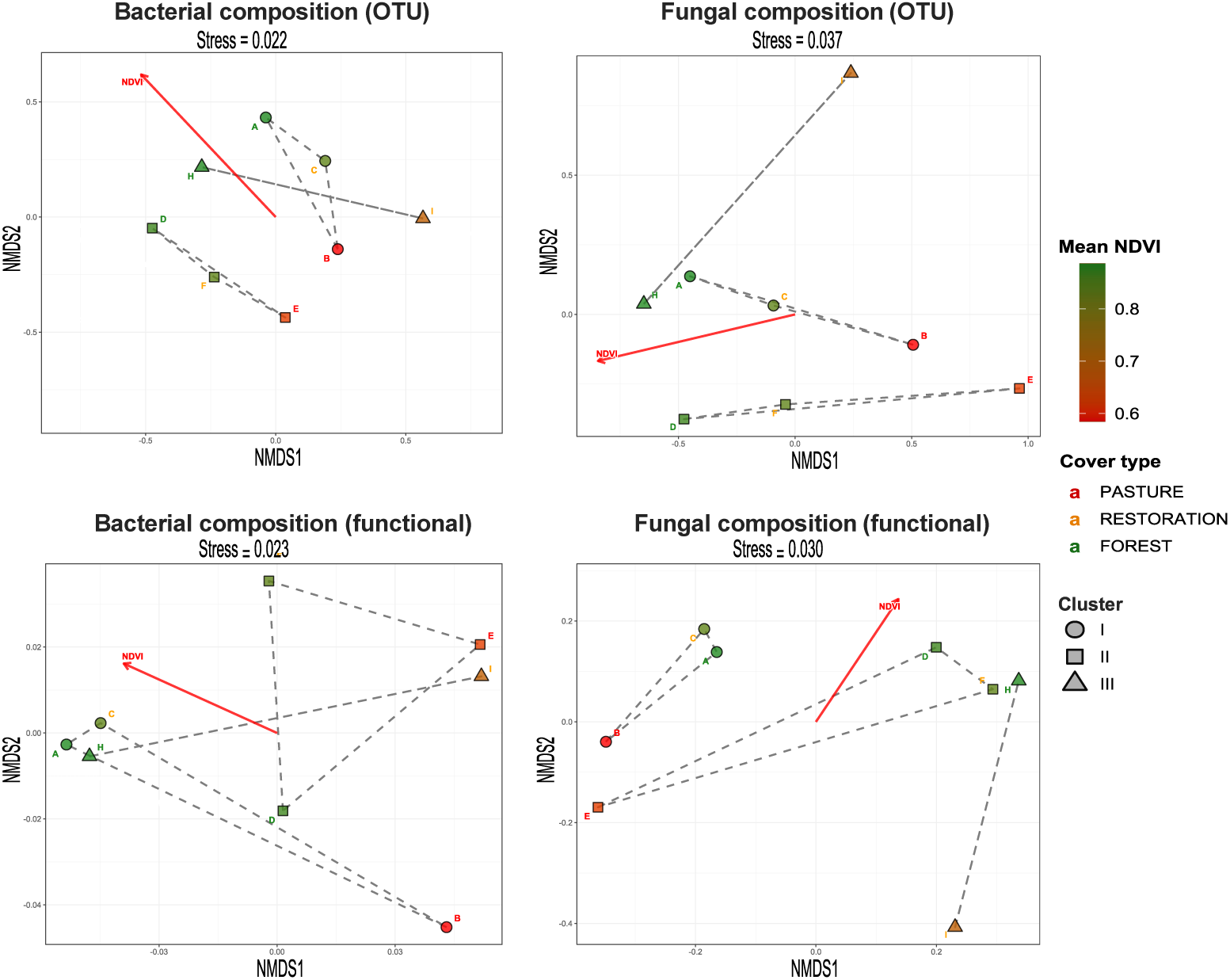
NMDS plots for OTU-based (top) and function-based (bottom) microbial composition. Red arrows represent the *cluster* and *NDVI* vectors when significant. Dashed lines connect samples from the same *cluster*.

PERMANOVA further confirmed the strong influence of both *NDVI* and *cluster* identity on microbial community composition (Table 2). Significant marginal effects were detected for *NDVI* in all datasets except in fungal functional guilds (although this was near-significant; p=0.067). *Cluster* identity did not show any significant marginal effect. All conditional tests showed a significant effect of both *cluster* and *NDVI*.

**Table 2:**
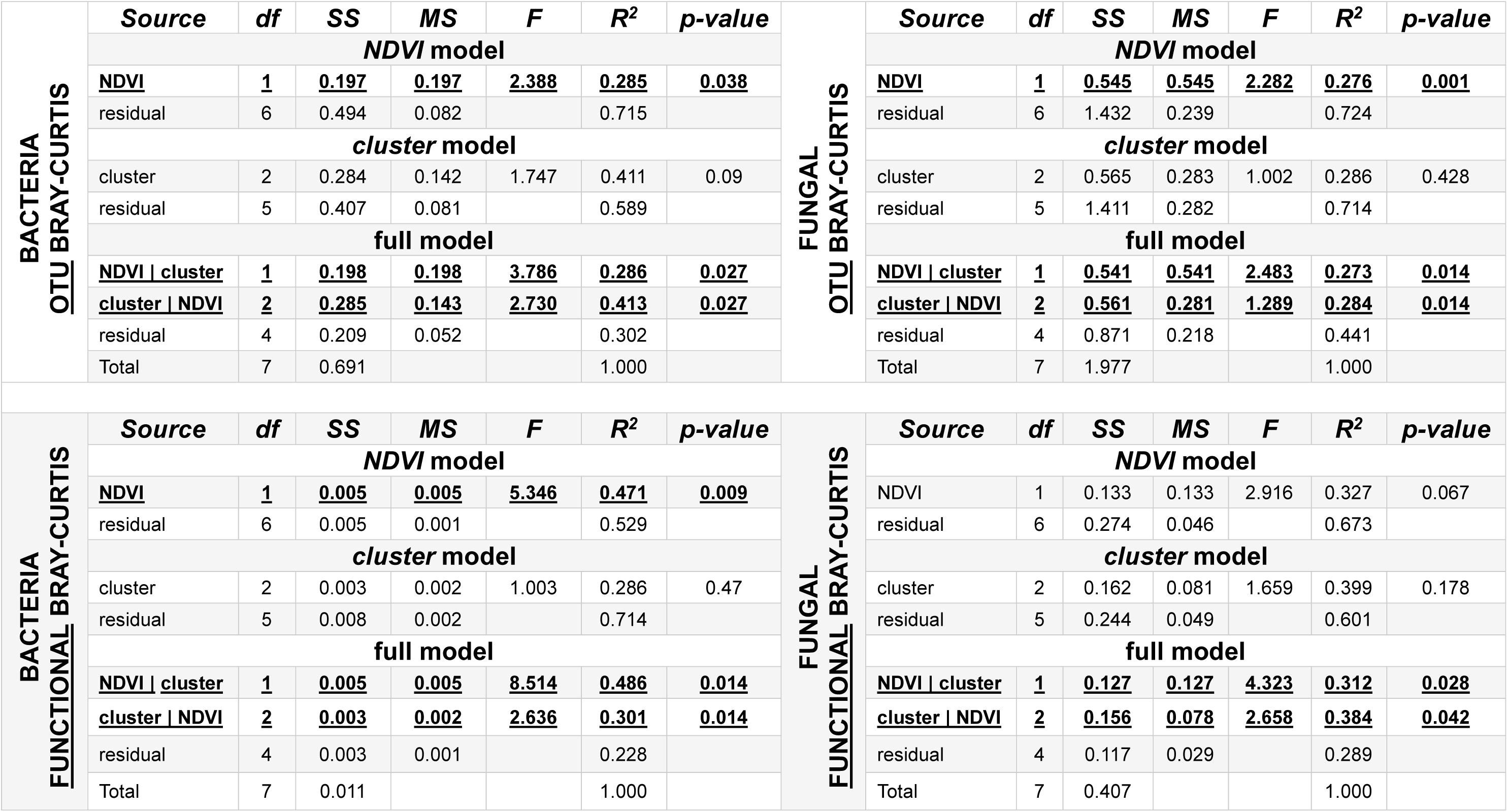
PERMANOVA results for OTU-based and function-based datasets (bacteria on left column, fungi on right). Note that similar p-values arose because of the low sample size, which established a ceiling on the number of permutations possible and thus the p-value intervals.

### Bacteria versus fungus diversity

There was no significant correlation (Spearman) between fungal and bacterial Shannon diversity.

However, the OTU-based Bray-Curtis dissimilarity matrices for bacteria and fungi were highly correlated (Mantel r: 0.690, p=0.0001; Figure 3). There was no correlation between the two function-based beta diversity matrices (r: 0.22, p=0.122).

**Figure 3:**
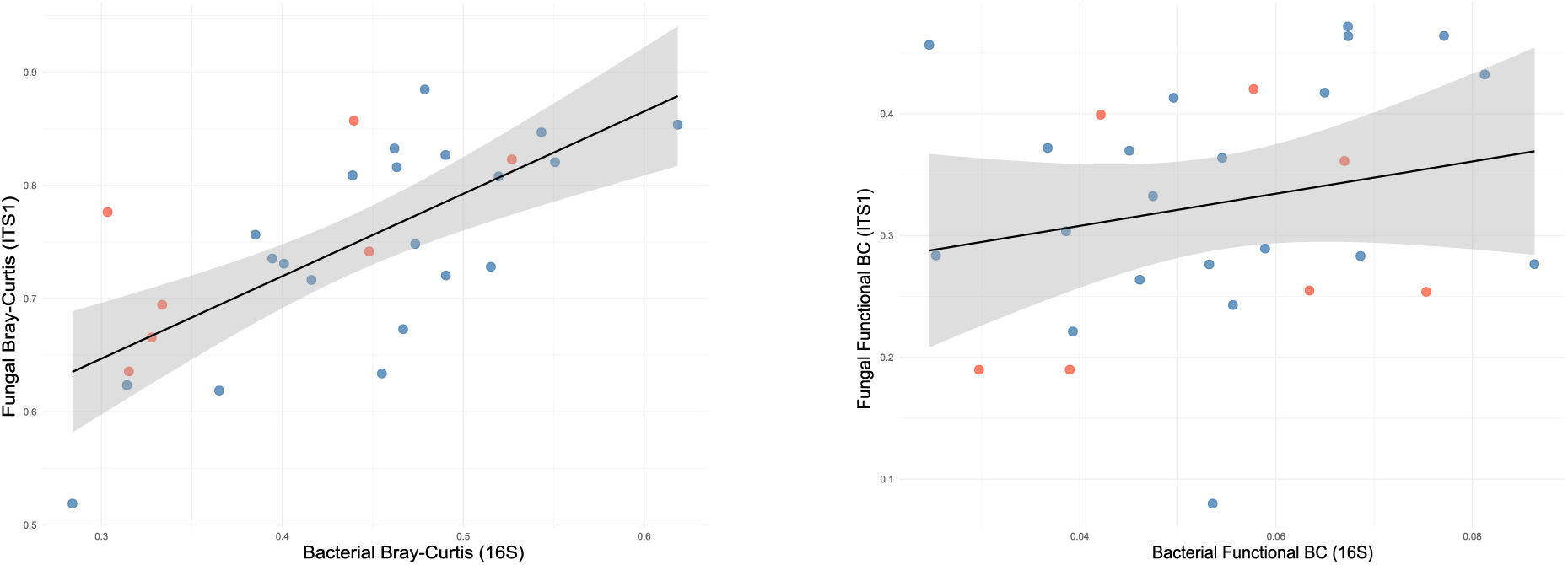
Correlation between Bray-Curtis dissimilarities of bacteria and fungi communities for *OTU*-based (left) and *function*-based matrices (right). Values of within-*cluster* comparisons are labelled in red, and between-cluster comparisons are in blue.

### Soil chemical profiles

In the soil chemistry dataset, 68.4% of total variance among sites was explained by the first two PCA axes. Graphically, samples grouped partly by geography (*cluster)*, especially *clusters I* and *II* (Figure 4).

**Figure 4:**
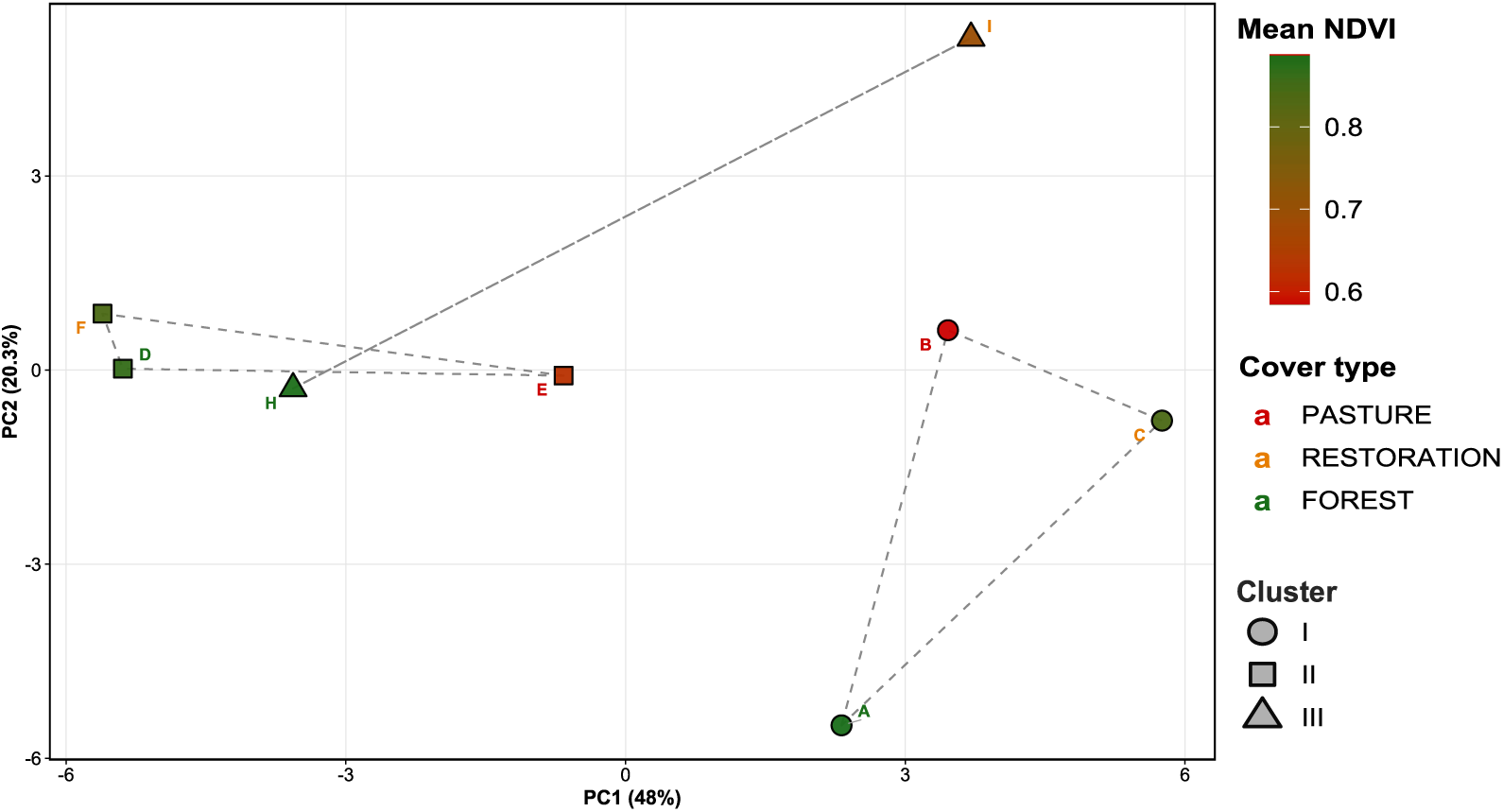
PCA ordination of 41 soil chemical parameters showing samples in relation to the first two components. Icon shape indicates *cluster* identity, color indicates *NDVI* values, with green being high vegetation density. Letter colors indicate if the sample was taken from a pasture, restored location, or reference forest. Dashed lines connect samples from the same geographic *cluster*.

Variance partitioning of the chemical profiles showed a significant joint-effect from *NDVI*+*cluster* (adj. R^2^ = 0.40, p =0.024; *SupplementalData.xlsx,* TAB *“Chemistry_varpart”*). This signal was primarily due to the contribution of *cluster* as, by itself, it had a near-significant marginal effect (adj. R^2^ = 0.29, p = 0.056) and after removing the contribution of *NDVI*, explained a substantial portion of the variance in chemical profiles (R^2^=0.37, p=0.03). There was no marginal or conditional effect of *NDVI* on chemical profiles.

## Discussion

### Alpha diversity

Neither bacterial nor fungal OTU Shannon diversity was significantly associated with *cluster* or *NDVI* (Table 1). However, bacterial *functional* Shannon diversity declined significantly with increasing *NDVI* (Adj. R^2^ = 0.749, p = 0.001; Table 1; Figure 1). This counterintuitive decline with increased vegetation may reflect bacterial niche contraction in mature forests, where increased recalcitrant carbon from woody litter selects for reduced functional assemblages compared to grassland/early successional soils (Zhou et al., 2017). Regardless of mechanism, decreasing bacterial functional Shannon diversity along the pasture–restoration–forest gradient may serve as a directional biomarker of successional advance in the Atlantic Forest.

Fungal functional Shannon diversity, by contrast, responded to marginal *cluster* rather than *NDVI* (Adj. R^2^ = 0.613, p = 0.049; Table 1), potentially indicating that edaphic conditions, which may be geographically structured given the spatial scope of our collections (see <u>Chemical profiles</u> section below), determine more fungal functional alpha diversity than the vegetation cover (=NDVI; Louisson et al., 2024).

### Beta diversity

Both bacterial and fungal community composition, assessed as Bray-Curtis dissimilarities in PERMANOVA, responded consistently to revegetation progression (i.e., *NDVI*) at both OTU and functional levels, and did so independently of *cluster* identity (Table 2). That *NDVI* explains meaningful variance in microbial composition beyond *cluster* identity confirms that the vegetation recovery signal represents a genuine, global, and biological response rather than purely a spatial/geographic artefact. This cross-site consistency provides empirical grounding for the use of soil eDNA metrics in operational restoration assessments (Toro et al., 2025).

Notably, the restored site in *cluster III* (labeled as “I” in Figure 2) generally plots closer to pasture samples in ordination space, which is explained because this site was substantially younger (∼2 years) than its restoration cohorts in *clusters I* and *II* (each >10 years). This pattern is visible in both bacterial and fungal NMDS plots and is consistent with restoration progress as a predictable driver of microbial succession toward the reference state.

The strong congruence between the *OTU* Bray-Curtis matrices (Mantel r = 0.690, p = 0.0001) suggests that, to some extent, both kingdoms share taxonomic drivers. However, a correlation between bacterial and fungal *functional* matrices was absent

### Chemical profiles

Soil chemistry was principally structured by geography, with *cluster* identity significantly explaining 29.1% of marginal variation and 37.3% when conditioned on *NDVI* (Figure 4; *SupplementalData.xlsx,* TAB *“Chemistry_varpart”*). The absence of any significant association between soil chemistry and *NDVI*, a proxy for vegetation density, indicates that edaphic conditions in the sampled landscape likely reflect historical land use rather than current vegetation status.

### Conclusions

Although our results are admittedly exploratory due to the limited sample sizes, they support the use of microbial eDNA to assess soil restoration trajectories, at least on the spatial scale sampled herein (∼40-km between the most distant clusters).

In terms of alpha diversity, restoration progress only manifests through functional Shannon diversity. In bacteria, this was only significant for *NDVI* (whether conditioned on *cluster* or not; Table 1). Restoration, as measured through *NDVI*, was not correlated to any fungal diversity metric, although there was a significant difference between *clusters* for functional diversity when conditioned on *NDVI*. Thus, functional alpha diversity, particularly for bacteria, may provide the more robust metric in any future framework for restoration trajectory monitoring.

The beta diversity analyses did not show a significant marginal effect of *cluster* on Bray-Curtis diversity for either microbe (Table 2). Only after conditioning on *NDVI* did *cluster* membership become significant. This dynamic may be partly driven by soil chemistry, whose variance distribution was significantly localized (i.e. associated with *cluster* identity), regardless of successional status of samples (*SupplementalData.xlsx,* TAB *“Chemistry_varpart”*). Importantly, this implies that chemical indicators alone are not reliable proxies of restoration trajectory because site legacy and spatial heterogeneity of soil can obscure early successional signals, at least over the decadal timescales evaluated herein.

Conversely to *cluster*, marginal and conditional *NDVI* effects were almost always significant for both kingdoms and for both OTU and functional levels. This suggests that, although *cluster* (=distance) does play a role in explaining DNA beta diversity, when only a raw metabarcoding dataset is available, a substantial portion of its variance may explain the restoration trajectory without consideration to distance of reference forest (at least at the spatial scale we evaluated).

That soil microbial communities in the Atlantic Forest retain a faithful and spatially scalable representation of restoration trajectories is particularly relevant for operational reference-site benchmarking, where site managers may lack an immediately adjacent reference forest. Such benchmarking will be substantially more robust with increased sampling density and when year-on-year measurements are taken. The latter will presumably maximize the metabarcoding signal and mitigate the confounding role of regional edaphic variations.

## Supporting information

Sites.kml

## Acknowledgments

Funding was kindly provided by the AstraZeneca Forest reforestation program and the Whitley Fund for Nature. The permit to access genetic data (Comprovante de Cadastro de Acesso ao Patrimonio Genético) was # AED7686. HBG received a CAPES scholarship (88887.677298/2022-00). We greatly appreciate the access given to us by the proprietors and residents of Fazenda Rosanela, Fazenda Estrela, and Fazenda San Maria and the extensive support provided by Alejandro at *Bioresearch do Brasil*.

## Data Availability Statement

Sequence data are deposited in NCBI SRA BioProject PRJNA1442186.

